# REPIC — an ensemble learning methodology for cryo-EM particle picking

**DOI:** 10.1101/2023.05.13.540636

**Authors:** Christopher JF Cameron, Sebastian JH Seager, Fred J Sigworth, Hemant D Tagare, Mark B Gerstein

## Abstract

Cryo-EM (cryogenic electron microscopy) particle identification from micrographs (i.e., picking) is challenging due to the low signal-to-noise ratio and lack of ground truth for particle locations. Moreover, current computational methods (“pickers”) identify different particle sets, complicating the selection of the best-suited picker for a protein of interest. Here, we present REPIC, an ensemble learning methodology that uses multiple pickers to find consensus particles. REPIC identifies consensus particles by framing its task as a graph problem and using integer linear programming to select particles. REPIC picks high-quality particles when the best picker is not known a priori and for known difficult-to-pick particles (e.g., TRPV1). Reconstructions using consensus particles achieve resolutions comparable to those from particles picked by experts, without the need for downstream particle filtering. Overall, our results show REPIC requires minimal (often no) manual picking and significantly reduces the burden on cryo-EM users for picker selection and particle picking.

**Availability:** https://github.com/ccameron/REPIC

---

Cryogenic electron microscopy (cryo-EM) (1) is a modern biophysical technique for protein structure determination. Protein complexes in solution are frozen and then imaged with electrons to produce various 2D projections (i.e., particles) within a digital electron micrograph. Individual particles in a micrograph are selected (i.e., picked), and then computationally registered to produce 3D reconstructions of the imaged protein complex. Protein crystallization is not required before cryo-EM imaging, and complexes can theoretically be as small as 17 kDa (2). However, micrographs have a low signal-to-noise ratio (SNR) due to limited electron beam exposure before proteins are damaged (3). To overcome low SNR, cryo-EM studies require hundreds to thousands of micrographs (4) from which as many as millions of particle images are selected. These datasets range in size from hundreds of gigabytes to several terabytes (5), with modern microscopes generating 10-20 terabytes a day (6).

Identifying particle images in a micrograph (i.e., particle picking) is a major bottleneck for cryo-EM image processing because of low SNR, sample contamination (e.g., ice crystals), and image artifacts in micrographs. Manually picking particles is impractical given the large number of micrographs. Computational methods, called particle pickers, including reference/template-matching (7–17) or machine learning algorithms (typically convolutional artificial neural network [CNN] based) (18–35), have been developed to automate particle picking. Currently available pickers typically come pre-trained on a large dataset (including both real and simulated data, which we refer to as “out-of-the-box”) or have the option to be initialized and retrained on a new set of micrographs (*ab initio*). Due to a lack of ground truth for cryo-EM particle locations (36), picker training is traditionally based on manually picked particles.

Current pickers have substantial error rates, requiring typically 50% (and often 80%) of their picked particles to be removed downstream by manual selection of particle clusters. Furthermore, these clusters are obtained through 2D or 3D classification and are not reproducible across cryo-EM pipelines (EMAN2 (10), CryoSPARC (16), RELION (37), etc.). In addition, each picker picks a different particle set due to its individual particle-background decision boundary. Differences in decision boundaries arise from each picker’s algorithm and training data. Since no single picker works best for all proteins and datasets, researchers rely on downstream processing to evaluate the quality of picking. This approach has real-world, practical limitations: First, choosing which 2D or 3D classes to keep or exclude is a subjective decision that requires sophisticated judgment. Second, a significant amount of manual picking is essential to train pickers, and this process also involves making subjective choices. Overcoming these limitations requires significant experience with cryo-EM image processing, which may not be available to all users. Therefore, more reliable automatic picking is of great interest in removing user bias and accelerating cryo-EM image processing.

One solution to this problem is to employ multiple pickers and use an algorithm to reconcile the different picker outputs without 3D reconstruction or manual interaction. One such approach has previously been reported by Sanchez-Garci et al. (2018) (24) called DeepConsensus (DC). DC iteratively combines picker output to produce intersection and symmetric difference particle sets. A separate CNN is then trained to pick particles using both particle sets as training data. While promising, the iterative creation of the intersection set results in a greedy algorithm and prevents global optimization. Further, intersection sets contain many false positives as particle bounding boxes densely overlap. Finally, training a separate CNN has the potential to introduce false positives and false negatives when the picker fits poorly to training data.

Here, we report an ensemble learning approach to the above problem, called REliable PIcking by Consensus (REPIC), that reconciles the output of multiple pickers into a consensus set of high quality particles. Ensemble learning is a machine learning technique that combines multiple algorithms to improve performance (38). As shown in Results, REPIC consensus particles produce high-quality reconstructions without 2D or 3D classification. REPIC can be used iteratively to *ab-initio* train pickers and produce high-resolution reconstructions for difficult-to-pick particles, requiring very few (1-3 particles per micrograph) or often no manual picking. Finally, REPIC allows for transfer learning between cryo-EM datasets, meaning that if manual picking is not possible for a target dataset, pickers can be trained using a closely related dataset. With REPIC, high-quality particles can be reliably picked, resulting in high-resolution reconstructions that are on par or superior to published results, without the need for additional particle filtering.

## Results

### REPIC algorithm

REPIC finds consensus particles from *k* picked particle sets in three steps (Figure 1A) by:

**Fig. 1.**
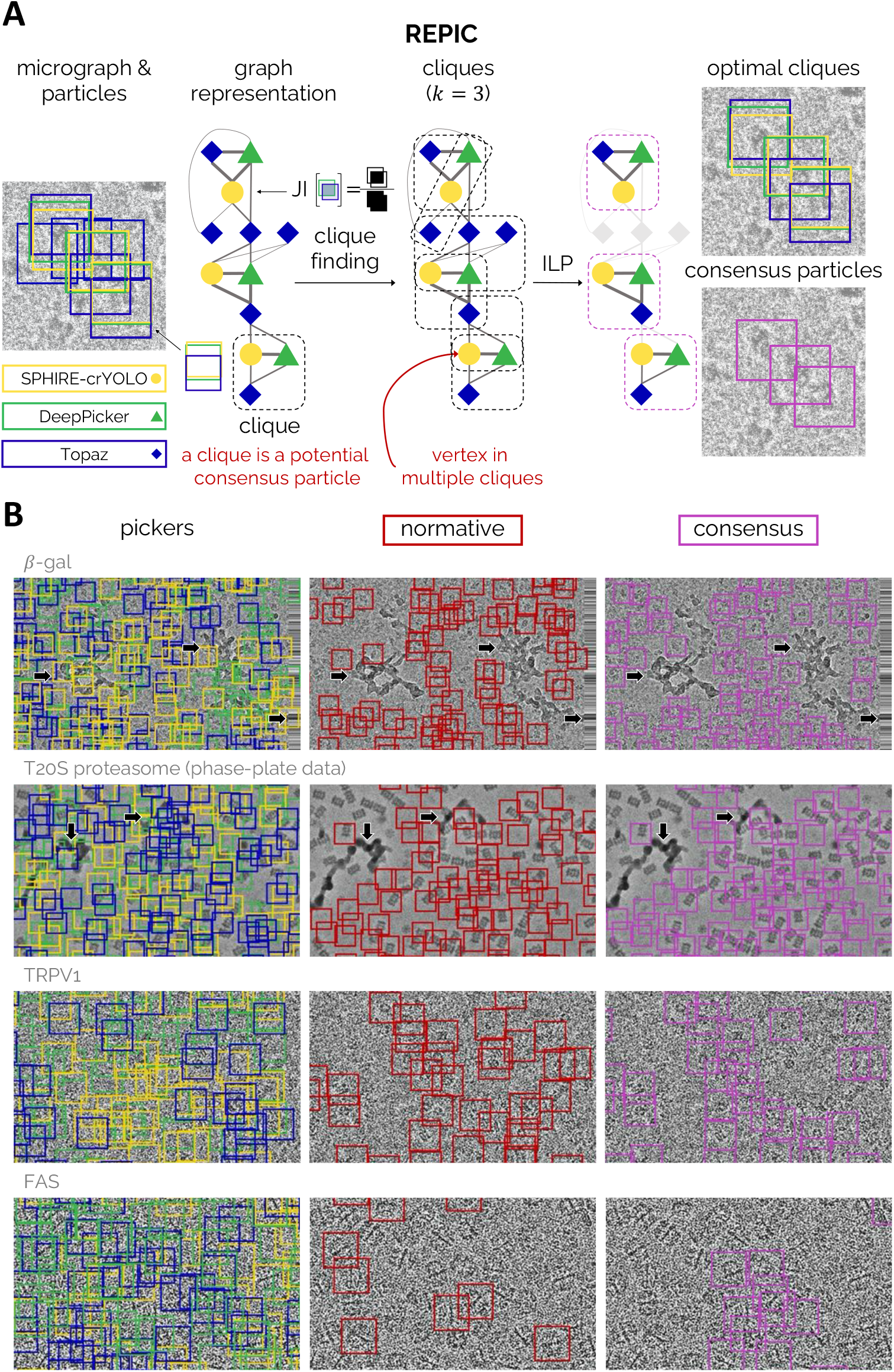
Identifying consensus particles with REPIC. **A**) Schematic representation of consensus particle identification by REPIC. Particle bounding boxes by individual pickers (SPHIRE-crYOLO (29) [yellow], DeepPicker (19) [green], and Topaz (26) [blue]) are represented as vertices in a computational graph. Edge weights are the overlap between two bounding boxes calculated by the Jaccard Index (JI). Clique finding is then performed, and an optimal subset of cliques is selected by integer linear programming (ILP — Supplemental Figure S1). Consensus particles are then derived from optimal cliques (see Online Methods). **B**) A random sampling of out-of-the-box picker (*left*), normative (red — *middle*), and consensus (purple — *right*) particles for (*top-to-bottom*) *—*-gal (EMPIAR-10017), T20S proteasome (10057), TRPV1 (10005), and FAS (10454) datasets. Arrows indicate sample contamination and image artifacts present in micrographs, which both normative and consensus particle sets were shown to avoid. TRPV1 and FAS micrographs have been low-pass filtered to make particles more visible.

**Fig. 2.**
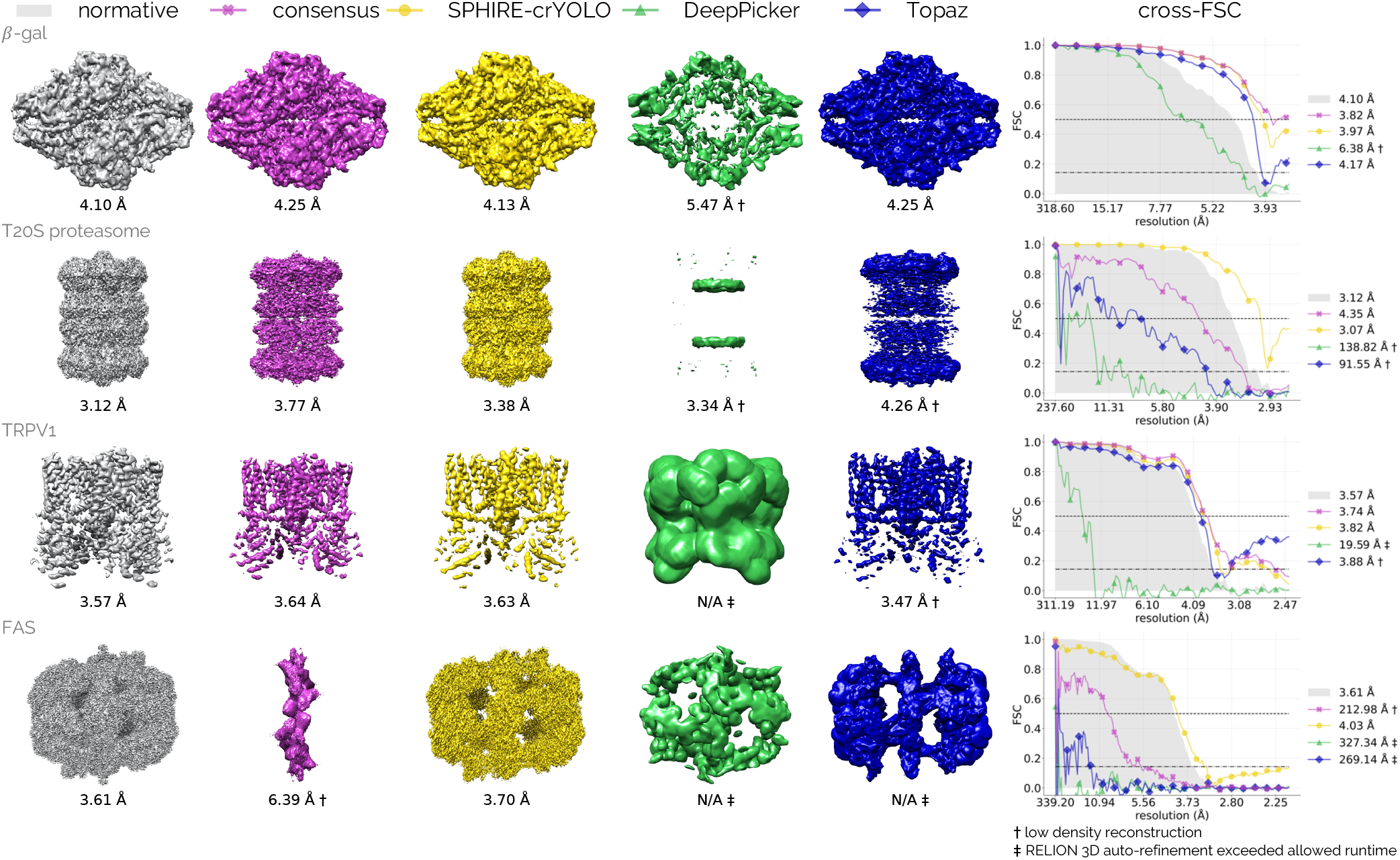
One-shot REPIC using out-of-the-box pickers. Out-of-the-box SPHIRE-crYOLO (yellow), DeepPicker (green), and Topaz (blue) pickers applied to (*top-to-bottom*) *—*-gal (EMPIAR-10017); in-focus, phase-plate T20S proteasome (10057); TRPV1 (10005); and FAS (10454) datasets. Consensus (purple) particles are identified from picker output using one-shot REPIC and shown to produce high-resolution densities when most pickers perform well. Cross Fourier shell correlation (cross-FSC) curves (see Online Methods) comparing picker and consensus maps to the normative (grey) are shown on the *right*. Normative curve is a half map FSC, while all other curves are cross-FSCs (see Supplemental Figure S3 for picker and consensus half map FSC curves). SPHIRE-crYOLO produces the highest-resolution, most-complete reconstructions for each dataset. Consensus particle sets represent the performance of the three-picker ensemble and obtain reconstructions comparable to the normative for all proteins except for the FAS dataset.

1. *Graph building* — representing particle bounding boxes as vertices in a computational graph. There is an edge between two vertices if the corresponding bounding boxes have a significant overlap as measured by the Jaccard Index. Edges only exist between bounding boxes of different pickers
2. *Clique finding* — identifying *k*-tuples of bounding boxes that have significant overlap with each other by finding cliques of size *k* in the graph
3. *Clique optimization* — selecting the subset of cliques with the maximum bounding box overlap and picker confidence subject to the constraint that each vertex participates in only one clique. Selected cliques represent consensus particles. Optimally selecting cliques is a combinatorial optimization problem; bounding boxes often overlap in a dense way so that a globally optimal grouping is not obvious (see Figure 1B *left*). REPIC uses integer linear programming (ILP) to obtain a non-greedy solution for clique selection (see Online Methods for more information)

REPIC makes minimal assumptions: It assumes that there are *k* pickers, and that all pickers provide a bounding box and score for each picked particle. The score takes values in [0, 1] and reflects a picker’s confidence in the particle. REPIC results reported below use three (*k* = 3) CNN-based pickers: SPHIRE-crYOLO (29), DeepPicker (19), and Topaz (26). SPHIRE-crYOLO and Topaz are modern (and widely accepted) pickers, while DeepPicker is an older picker. However, REPIC is not limited to CNN-based pickers and can be used with *k*-many pickers.

REPIC was tested using two modes:

1. *one shot* — REPIC takes the output of (possibly trained) pickers and finds high-quality consensus particles without 3D reconstruction using the three steps above. This mode relies on individual pickers being well trained on large datasets (Figures 1B and 2)
2. *iterative* — Taking inspiration from the work of McSweeney et al. (2020) (39): pickers are *ab-initio* trained using either one-shot REPIC output or manually picked particles. Pickers are run and one-shot REPIC is used to find consensus particles to retrain pickers. This pick-REPIC-retrain loop (Figure 3A) is then executed for a user-defined number of iterations (see Pseudocode 1)

For cases where out-of-the-box pickers may fail, we show that REPIC’s iterative mode improves picker performance using one of the following three initializations:

**Fig. 3.**
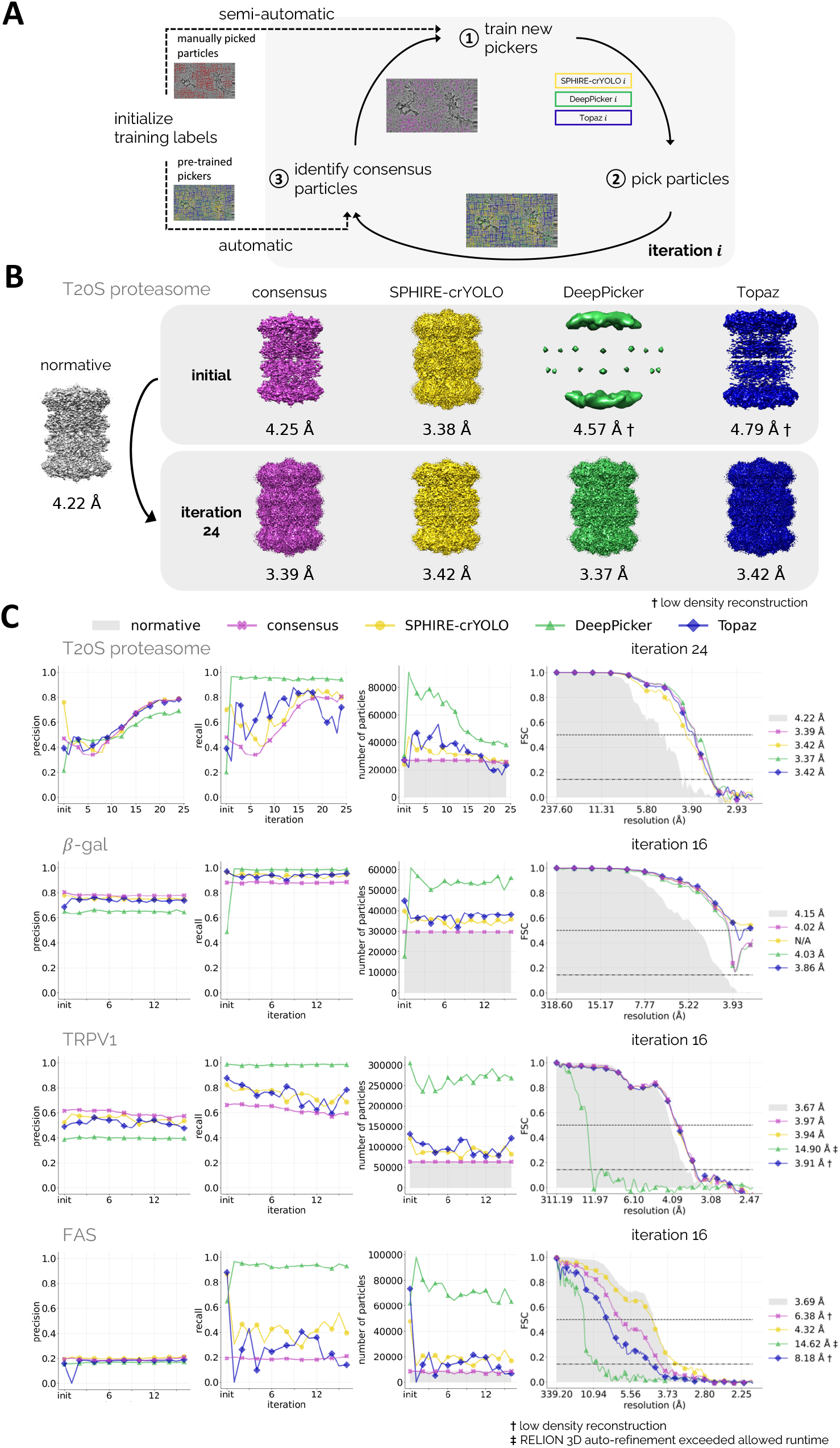
Iterative-mode REPIC initialized using out-of-the-box picker output. **A**) Schematic representation of REPIC’s iterative mode. Training labels (i.e., particles) can be provided by either pre-trained pickers (automatic) or manual particle picking (semi-automatic). New pickers are *ab-initio* trained from the training data and then used to pick particles. Consensus particles are identified using REPIC and used as training labels in the proceeding iteration. The pick-REPIC-train loop is then repeated for a user-defined number of iterations. A cross-validation strategy (see Online Methods) is used to subset micrographs into training, validation, and testing sets for picker training. **B**) T20S proteasome reconstructions obtained from normative (grey), consensus (purple), SPHIRE-crYOLO (yellow), DeepPicker (green), and Topaz (blue) picked particle sets from the initial (out-of-the-box picker output — *top*) and final (24 — *bottom*) iteration. All algorithms converge to a similar particle set (due to consensus particles being used as training labels) and achieve high-resolution densities in the final iteration. Most final pickers improve upon the initial, out-of-the-box pickers and outperform the normative particle set. **C**) Evaluation metrics (precision, recall, and number of particles) for iterative-mode REPIC initialized using out-of-the-box picker output applied to (*top-to-bottom*) T20S proteasome (EMPIAR-10057), *—*-gal (10017), TRPV1 (10005), and FAS (10454) datasets. cross-FSC curves of reconstructions obtained from the final iteration are shown on the *right*. The same FSC resolution thresholds are used as described in Figure 2. Most algorithms show improvement over out-of-the-box pickers (Figure 2B) and produce similar final densities (Supplemental Figure S4 — except for DeepPicker and the FAS dataset).

I. out-of-the-box picker output (Figure 3)
II. manually picked particles (Figure 4)
III. *ab-initio* transfer learning (Figure 5)

### REPIC evaluation

In place of ground truth, REPIC was evaluated using four published particle sets obtained from the EMPIAR resource (https://www.ebi.ac.uk/empiar/): TRPV1 (EMPIAR-10005), *—*-galactosidase (*—*-gal — 10017), T20S proteasome (10057), and fatty acid synthase (FAS — 10454). These particle sets contain high-quality particles selected by downstream image processing and represent the norm or expected particle set. We refer to these sets as the “normative”. For example, only 111, 000 particles of the original 857, 000 particles are found in the FAS normative set (before random selection of micrographs — see Online Methods and Supplemental Data File 1). When possible, the normative particle set consists of the particle set that produced the final, published reconstruction (see Online Methods for more information). All datasets, except T20S proteasome, were acquired with standard defocus-contrast imaging. T20S proteasome micrographs were captured in-focus using a phase plate.

**Fig. 4.**
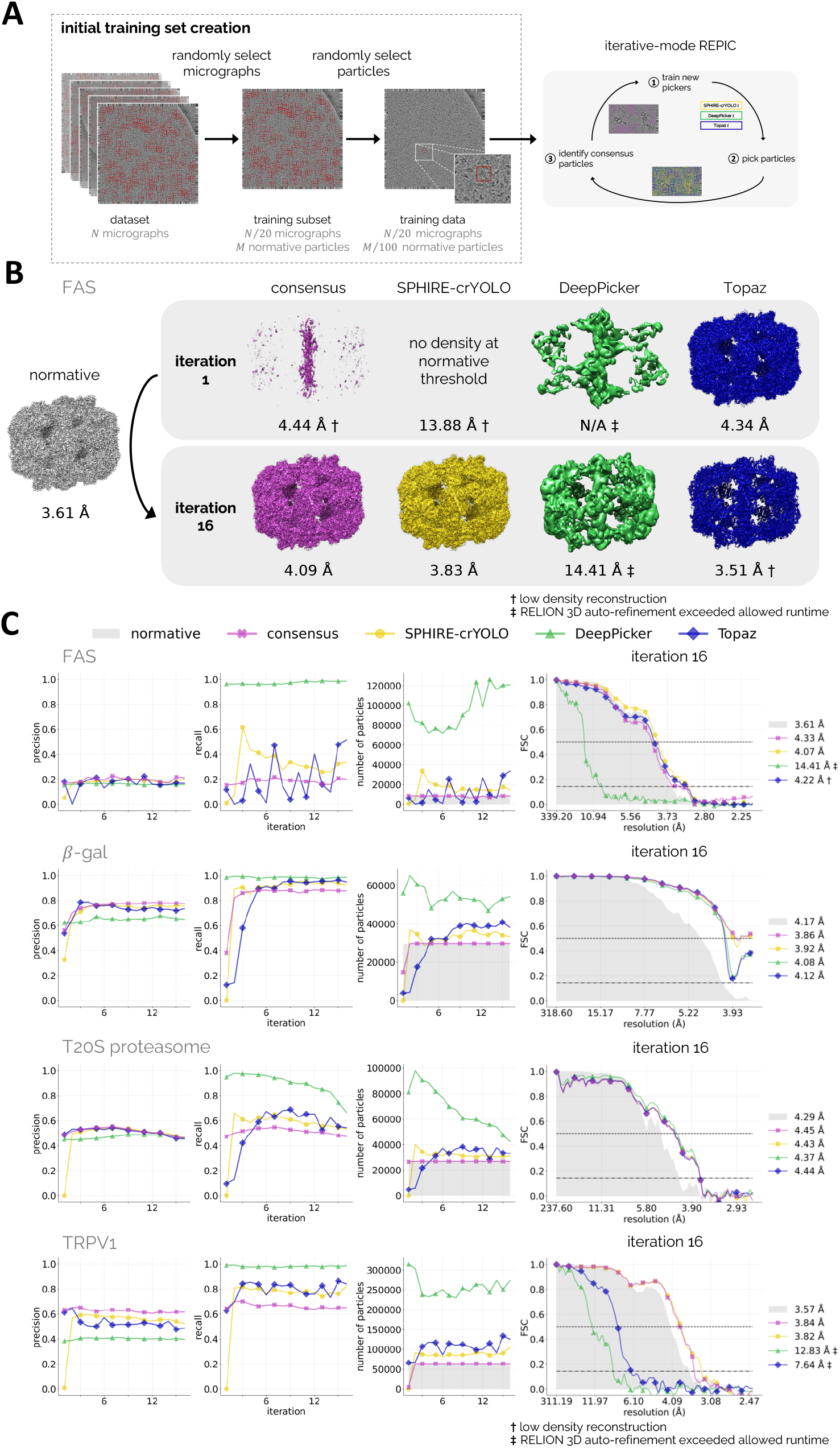
Iterative-mode REPIC using minimal manual picking for *ab-initio* picker training. **A**) Schematic of the three step, random selection analysis applied to each cryo-EM dataset: (1) 5% of micrographs are randomly selected as the training subset, (2) 1% of normative particle coordinates are randomly selected and chosen as training labels, and (3) training labels and micrographs are provided to REPIC’s iterative mode. See Online Methods for more information. **B**) FAS resonstructions obtained from normative (grey), consensus (purple), SPHIRE-crYOLO (yellow), DeepPicker (green), and Topaz (blue) picked particle sets from the initial (pre-trained picker output — *top*) and final (16 — *bottom*) iteration. All algorithms (except DeepPicker) converge to a similar, high-resolution reconstructions in the final iteration. **C**) Evaluation metrics (precision, recall, and number of particles) for iterative-mode REPIC of (*top-to-bottom*) FAS (EMPIAR-10454), *—*-gal (10017), T20S proteasome (10057), and TRPV1 (10005) datasets. Cross-FSC curves and resolutions for densities obtained from the final iteration are shown on the *right*. The same resolution thresholds are used as described in Figure 2. Most algorithms produce similar final, high-resolution densities (Supplemental Figure S5).

**Fig. 5.**
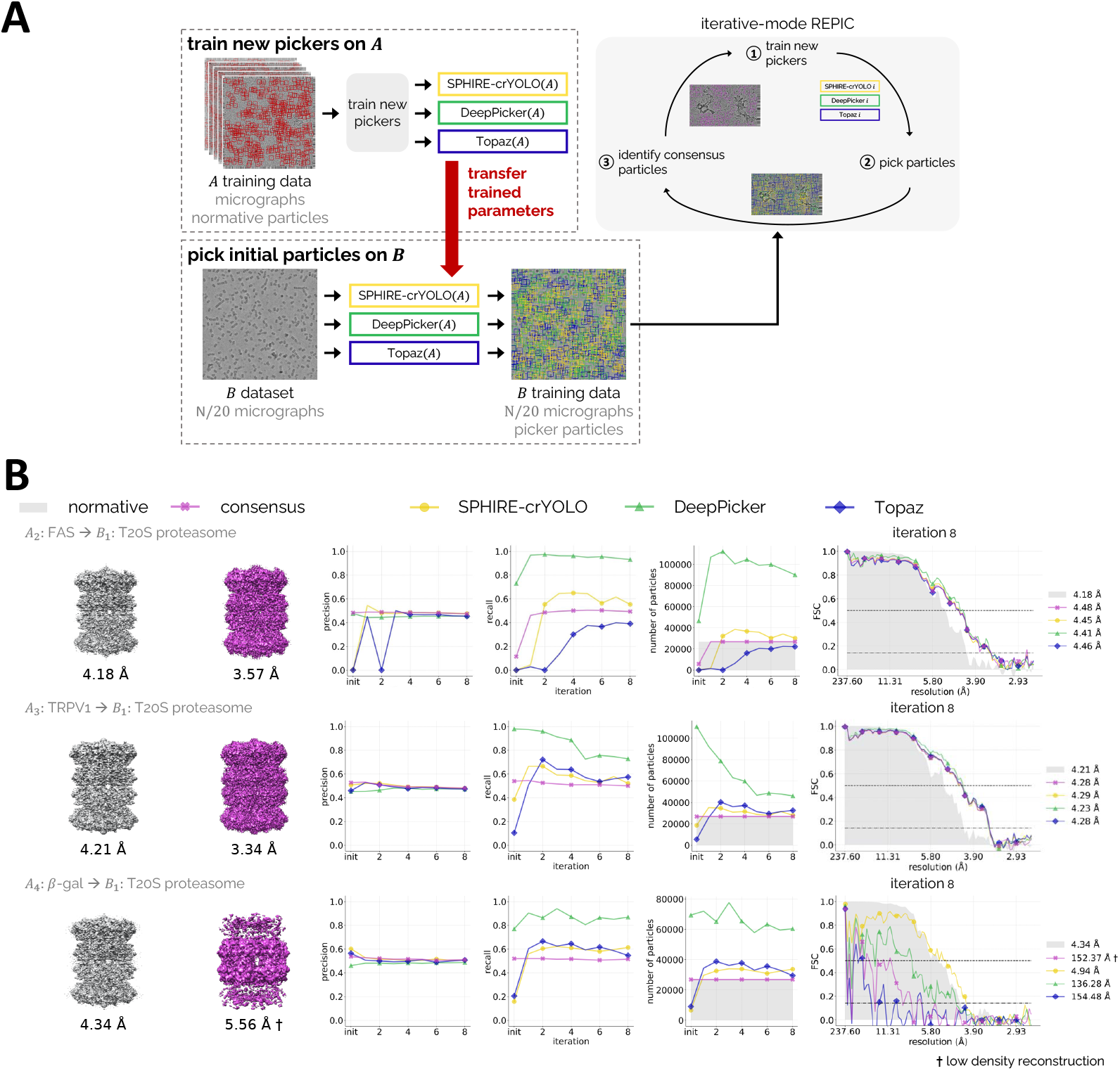
*Ab-initio* transfer learning using iterative-mode REPIC. **A**) Schematic representation of *ab-initio* transfer learning with REPIC: A set of SPHIRE-crYOLO, DeepPicker, and Topaz pickers are initialized and trained separately on cryo-EM dataset *A*. These pre-trained pickers are then applied to cryo-EM dataset *B* to generate initial training labels for iterative-mode REPIC. **B**) Normative (grey) and consensus (purple) reconstructions for T20S proteasome dataset resulting from REPIC’s iterative mode using pickers pre-trained on either the FAS (EMPIAR-10454 — *top*), TRPV1 (10005 — *middle*), and *—*-gal (10017 — *bottom*) dataset. Precision, recall, and number of (picked) particles curves are provided along with the cross-FSC (*right-most column* — using thresholds described in Figure 2). High-resolution reconstructions that improve upon the normative particle coordinates are achieved using FAS and TRPV1 datasets to pre-train pickers. However, using *—*-gal to pre-train pickers fails to produce a useful density (most likely due to the pixel resolution difference of both datasets — see Supplemental Data File 2). *Ab-initio* transfer learning relies on the similarity of the training data and target cryo-EM dataset. When datasets are too dissimilar, this approach will fail to produce reconstructions (see Supplemental Figure S7).

Particle sets produced by a picker or REPIC were characterized by comparing to the normative particle set using precision and recall. These particle sets were also separately used to obtain reconstructions with RELION. All reconstructions were produced using the same procedure without 2D or 3D classification (see Online Methods for more information). Supplemental Data Files 2 and 3 list picker & RELION parameters and the number of picked particles for each particle set. Reconstructions from picked particle sets by individual algorithms (SPHIRE-crYOLO, DeepPicker, Topaz, and REPIC) are compared to the reconstruction obtained from the normative particle set using a cross Fourier shell correlation (cross-FSC — see Online Methods). Since the cross-FSC is derived from non-disjoint particle image sets, the cross-FSC resolution is reported at FSC= 0.50. Cross-FSC curves indicate the resolution at which the similarity between the normative reconstruction and a consensus or picker reconstruction drops below 50%.

### One-shot picking

To show REPIC can find useful consensus particle sets even when the pickers have performance variation, out-of-the-box pickers were used and one-shot REPIC was applied to find consensus particles. In all datasets except the T20S proteasome, REPIC consensus particles have the highest precision (Table 1), which supports an ensemble learning approach to particle picking.

**Table 1.** Precision (prec), recall (rec), and number (num) of picked particles for one-shot REPIC with out-of-the-box pickers. The number of normative particles is listed in parentheses under each dataset. Picker particle sets are filtered for false positives (see Supplemental Information).

| dataset | consensus |  |  | SPHIRE-crYOLO |  |  | DeepPicker |  |  | Topaz |  |  |
| --- | --- | --- | --- | --- | --- | --- | --- | --- | --- | --- | --- | --- |
|  | prec | rec | num | prec | rec | num | prec | rec | num | prec | rec | num |
| β-gal<br>(40,863) | **0.812** | 0.883 | 40,824 | 0.788 | 0.968 | 39,838 | 0.660 | 0.498 | 17,741 | 0.695 | **0.972** | 44,680 |
| T20S proteasome<br>(35,469) | 0.470 | 0.477 | 35,392 | **0.759** | **0.690** | 23,975 | 0.215 | 0.200 | 30,310 | 0.393 | 0.388 | 27,047 |
| TRPV1<br>(80,443) | **0.616** | 0.659 | 80,655 | 0.542 | 0.822 | 120,888 | 0.390 | **0.989** | 305,425 | 0.489 | 0.877 | 131,402 |
| FAS<br>(11,390) | **0.196** | 0.192 | 11,040 | 0.194 | 0.868 | 47,824 | 0.155 | 0.642 | 61,877 | 0.158 | **0.881** | 73,167 |

SPHIRE-crYOLO picked high quality particles for all datasets (based on the resulting reconstructions and cross-FSC resolutions — Figure 2), while Topaz picked good quality particles for three of the four datasets (FAS was the exception). DeepPicker picked moderate quality particles for *—*-gal and TRPV1, and poor quality particles for the proteasome and FAS datasets (RELION failed to finish within the allowed runtime for the proteasome dataset — indicated by *‡*, see Online Methods for more information). REPIC consensus particles achieved high-resolution reconstructions except for FAS where two of the three pickers failed. Similar behaviour is also observed with RELION 2D classes (Supplemental Figure S2). These results show that when one-shot REPIC is used, the consensus particle set is consistent with the best picker, <u>even when the best picker is not known a priori</u>.

### Iterative ensemble particle picking

To demonstrate how REPIC can be used on a novel particle, REPIC’s iterative mode (Figure 3A) was initialized using out-of-the-box picker output and pickers were then *ab-initio* trained. REPIC was separately applied to the four cryo-EM datasets for 16 iterations. The T20S proteasome dataset was processed for an additional eight iterations to observe a plateau in its precision curves. Figure 3B highlights the significant improvement that can be achieved between the initial and final consensus particle sets for the T20S proteasome dataset. DeepPicker showed the largest improvement, where the achieved reconstruction improves from an erroneous, disconnected map (4.57 Å) to a high-resolution map (3.37 Å). All four algorithms achieve reconstructions that are almost an Angstrom higher in resolution compared to normative particle coordinates (3.37-.42 vs. 4.22 Å).

Figure 3C (and Supplemental Figure S4) shows the results of REPIC’s iterative mode initialized with out-of-the-box picker output for all four tested cryo-EM datasets. REPIC processing of the T20S proteasome dataset demonstrates how the iterative mode can improve both precision and recall over multiple iterations. Convergence rates of the iterative algorithm vary from particle to particle and the final particle set may be dissimilar from the normative. For example, REPIC processing of the *—*-gal dataset shows how the picker ensemble may converge during early iterations and have precision and recall remain stable across later iterations (Figure 3C). TRPV1 and FAS datasets demonstrate that the framework may converge to a picked particle set that is dissimilar (to different degrees) from the normative particle set while achieving high-resolution reconstructions (see Supplemental Figure S4 for final iteration reconstructions). DeepPicker fails to achieve high-resolution reconstructions for the TRPV1 and FAS datasets, which is most likely due to the large number of particles picked by this picker even after false positive filtering. The precision and recall of the consensus particle set remains consistent across iterations, even when individual pickers (e.g., Topaz) show instability (large changes in precision and/or recall). In fact, the precision of the consensus particle set remains as high or higher than any individual picker. The cross-FSC curves and resolutions by consensus maps are either improved (e.g., from an erroneous reconstruction to improved FAS reconstruction at 4.47 Å) or remain consistent (*—*-gal reconstruction at ∼4.3 Å) compared to the out-of-the-box pickers.

#### Semi-automatic picking by iterative ensemble

Next, we asked if high-resolution reconstructions could be obtained using iterative-mode REPIC and a minimal, initial particle set. For each cryo-EM dataset, 5% of micrographs were randomly selected as the training subset (Figure 4A).

For each micrograph in the training subset, 1% of normative particle coordinates were randomly selected to represent manually picked particles and *ab-initio* train pickers. In total, 6, 6, 8 and 44 training particles were selected for the *—*-gal, T20S proteasome, FAS, and TRPV1 datasets, respectively. The TRPV1 initial particle set contains a factor of six or more particles due to the number of training micrographs selected, which is based on the total number of micrographs (*N* = 849 — see Supplemental Data File 2 for more information). Initial training labels and micrographs were then provided to iterative-mode REPIC and 16 iterations were performed.

Figure 4B shows significant improvement in FAS reconstructions between the first and last iteration using REPIC’s iterative mode. In the first iteration, the consensus particle set is unable to produce a reconstruction due to the poor performance of SPHIRE-crYOLO and DeepPicker (based on their respective reconstructions). After 16 iterations, the iterative framework produces both consensus and SPHIRE-crYOLO reconstructions at a resolution approaching the normative. These high-resolution reconstructions are achievable because the ensemble is composed of multiple pickers of different architectures and objective functions. Pickers that require minimal training data drive the initial iterations of iterative-mode REPIC (based on the resulting reconstructions). Later iterations are driven by pickers able to achieve higher-resolution reconstructions.

Initializing iterative-mode REPIC with a minimal particle set results in high-resolution reconstructions on par with (*—*-gal and TRPV1 datasets), approaching (FAS), and better than (T20S proteasome) the normative particle set (Supplemental Figure S5). *—*-gal, T20S proteasome, and TRPV1 precision and recall curves (Figure 4C) show how the ensemble quickly improves and converges over a small number of iterations (six or less) from a small subset of initial training labels. SPHIRE-crYOLO and Topaz show instability for the FAS dataset across iterations even though the final reconstructions are comparable to the normative map (based on RELION-estimated resolution and cross-FSC analysis — Figure 4C). Final DeepPicker particle sets are also improved (based on cross-FSC curves and obtained resolution) over automatic iterative picking even though low-resolution reconstructions are achieved for the TRPV1 and FAS datasets (Supplemental Figure S5). Similar to automatic runs of the iterative mode, the precision and recall of consensus particle sets remain stable across iterations and lead to high-resolution reconstructions that are comparable to the normative map.

#### Ab-initio transfer learning

Finally, we explored the potential for transfer learning between two individual cryo-EM datasets (*A* and *B*) using REPIC’s iterative mode (Figure 5A). Pickers are ab-initio trained using training data from dataset *A*. All micrographs and normative particles from dataset *A* (minus a held-out subset for validation — see Online Methods) were used to train pickers. Trained pickers are then applied to a training subset of micrographs from dataset *B* to generate the initial training labels for iterative-mode REPIC. Four sets of the three studied pickers were trained (*A*_1_, *A*_2_, *A*_3_, and *A*_4_), each set specific to one of the four studied cryo-EM datasets (T20S proteasome, FAS, TRPV1, and *—*-gal, respectively). Picker sets *A*_2_, *A*_3_, and *A*_4_ were then used to pick T20S proteasome particles (*B*_1_ — Figure 5B). Using pickers trained on the FAS and TRPV1 datasets resulted in high-resolution reconstructions that improved upon the achieved resolution of normative reconstruction. However, using the *—*-gal dataset to pre-train pickers led to only a reasonable reconstruction using SPHIRE-crYOLO picked particles, with all other algorithms obtaining lower-resolution reconstructions compared to the normative (Supplemental Figure S6). While promising, caution must be taken when performing transfer learning using REPIC’s iterative mode, as shown in Supplemental Figure S7, FAS (*B*_2_) reconstructions were not able to be produced.

## Discussion

In one-shot mode, REPIC produces high-quality consensus particle sets when multiple out-of-the-box pickers are used, and the identity of the best picker is not known. REPIC works reliably as long as a majority of the pickers perform reasonably well. If most pickers fail, then the notion of consensus particles is not very meaningful. Such is the case with the FAS dataset in Figure 2.

In the one-shot experiments, it is interesting to observe that SPHIRE-crYOLO obtained the highest precision and recall with the T20S proteasome dataset (Table 1). We believe this result is due to SPHIRE-crYOLO being the only out-of-the-box picker trained on similar phase-plate data (EMPIAR-10050) (40).

In all three initializations used with REPIC’s iterative mode (Figures 3-5), consensus particle sets reliably produced high-resolution reconstructions, with resolutions comparable to the normative. These resolutions were achieved without downstream filtering (such as 2D and 3D classification). 2D and 3D classes are not reproducible across cryo-EM pipelines and are sensitive to the classification parameter values, number of particle images, and number of micrographs provided. In addition, identifying correct classes relies on a subjective choice by a cryo-EM expert. REPIC further automates particle picking by provide quality particle sets that either negate the need for or better initialize particle filtering.

When using REPIC’s iterative mode, individual pickers converge to similar reconstructions. This result is due to the *ab-initio* training using consensus particles. A minimal amount of training data (1% of particles per training micrograph) is required to initialize REPIC’s iterative mode. Figure 4B illustrates the benefits of using a picker ensemble: REPIC, along with pickers, can bootstrap from picking poor particles in the early iterations to picking high quality particles in the later iterations (based on the resulting reconstructions). In Figure 4B, early iterations of REPIC are driven by Topaz, which requires less training data due to its positive-unlabeled learning algorithm. Later iterations are driven by SPHIRE-crYOLO which requires more training data but achieves higher resolution reconstructions.

The above results support the main claims of this paper: (1) ensemble learning, as manifested in REPIC, can provide high quality particles, even when the best picker is not known a priori; (2) in more difficult cases, minimal manual picking is sufficient to bootstrap REPIC into a high quality particle regime; (3) high-resolution reconstructions can be obtained with REPIC, without downstream processing such as 3D classification.

Future work will focus on improving the runtimes of both the one-shot and iterative modes of REPIC. Currently, an exhaustive search is performed when building the computational graph. A *k*-d tree approach could be used to significantly reduce the amount of bounding box comparison and improve REPIC runtime. During the iterative REPIC loop, pickers are run in sequence. *Ab-initio* training of pickers contributes to a significant amount of REPIC’s runtime (Supplemental Figure S8). Parallel application of pickers or a variant (i.e., only re-training the worst performing picker) can improve the speed of REPIC iterations.

## Online Methods

### Dataset description

Cryo-EM digital micrographs and normative particle coordinates were obtained from the Electron Microscopy Public Image Archive (EMPIAR) resource for entries EMPIAR-10005 (41), EMPIAR-10017 (37), EMPIAR-10057 (42), and EMPIAR-10454 (43). Associated, published 3D volumes for each EMPIAR dataset were retrieved from the Electron Microscopy Data Bank (EMDB) from entries EMD-5778, EMD-2824, EMD-3347, and EMD-4577, respectively. Due to the large number of micrographs in the EMPIAR-10454 dataset (*N* = 4, 593), 10% of micrograph and paired particle coordinate files (*N* = 460) were randomly selected and used in this study (see Supplemental Data File 1 for a list of the selected files). EMPIAR-10057 multi-frame micrographs were aligned and summed using MotionCor2 v1.5.0 (44) in RELION v3.1.3 (45). Contrast transfer function (CTF) estimation was performed for all datasets (except for the in-focus, phase-plate data of EMPIAR-10057) using CTFFIND4 v4.1.14 (46) in RELION. Please see Supplemental Data File 2 for a summary of RELION and CTFFIND4 parameters used to process each dataset.

Additional micrograph preprocessing was performed by each picker (i.e., low-pass filtering by SPHIRE-crYOLO (29), image standardization by DeepPicker (19), Gaussian mixture model [GMM] normalization by Topaz (26)) before particle picking. Picker installation, application, and false positive filtering are described in Supplemental Information.

### REPIC

REliable PIcking by Consensus (REPIC — /r ‘pik/) is a non-greedy approach to identifying consensus particles from *k* picked particle sets {*S*_1_,…, *S*_*k*_}. The input to REPIC is the set of picked particle sets 𝒮 = {*S*_1_,…, *S*_*k*_}, where each particle in a set is expected to have micrograph coordinates, a particle detection box size, and a quality score *s*. REPIC output is a consensus particle set in BOX file format. REPIC represents all picked particles in a micrograph as an undirected, *k*-partite graph *G* = (*V, E*). Each vertex *v* in *V* corresponds to a particle detection box from a picked particle set. Each edge *e* in *E* represents a pair of overlapping particle detection boxes. The edge weight *o* between two vertices is the overlap (i.e., the Jaccard Index) of their corresponding particle detection boxes. Intuitively, the goal of REPIC is to find the set of non-overlapping *k*-size cliques in *G* that maximize particle overlap and score. For each clique *c* = (*V*_*c*_, *E*_*c*_), the clique weight *w*_*c*_ is the product of its median edge weight 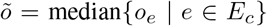 and median vertex score 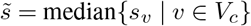. REPIC aims to find the disjoint set of cliques *𝒞* that maximizes ∑_*c*∈*𝒞*_ *w*_*c*_.

The details of the above steps are as follows:

1. *Graph building* — An undirected graph *G* is built from the output of *k* picked particle sets (as described above). In this study, picked particle sets are generated by three CNN-based pickers: SPHIRE-crYOLO (29), DeepPicker (19), and Topaz (26). Edges with *o <* 0.3 are considered to be particle detection boxes that do not overlap and are excluded from *G*.
2. *Clique finding* — All cliques of size *k* are enumerated using a modified Bron-Kerbosch algorithm (47), as implemented by the Python NetworkX package (48).
3. *Clique optimization* — A clique in the graph *G* corresponds to a single consensus particle. However each vertex in *G* (a picked particle) may participate in multiple cliques. To ensure each vertex associates with a single clique in the final set, cliques *x* are selected using Integer Linear Programming (ILP — Supplemental Figure S1) as follows: Suppose that the result of clique finding is *m* cliques containing *n* vertices. Define an *n × m* matrix *A*, where the element *A*_*ij*_ =1 is the *ith* vertex participating in the *jth* clique, else *A*_*ij*_ = 0. Then, the ILP is defined as below where *x*_*j*_ is a binary variable denoting whether the *j*th clique is selected (*x*_*j*_ = 1) or not (*x*_*j*_ = 0).

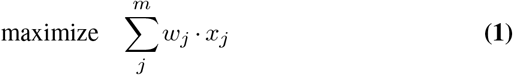

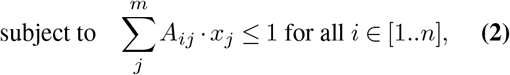

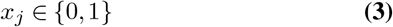

Equation 2 ensures vertices are only associated with a single clique by limiting row sums in *A* to 1. A globally optimal solution to the above problem is then found using ILP branch-and-bound optimization, as implemented by the Python Gurobi package (49).

Graph building, clique finding, and clique optimization are performed on a per-micrograph basis. REPIC itself does not require a GPU (although the pickers do) and runs efficiently on a single workstation with a processing time of seconds per micrograph. Graph building (specifically the exhaustive search for overlapping particle detection boxes) is the limiting step for REPIC (2-10 seconds per micrograph). The ILP solver is efficient (*<*0.2 seconds per micrograph).

#### One-shot mode

Given an initial set of picked particle sets *𝒮*_init_, one-shot REPIC executes the above steps once. In the discussion below, we denote the one-shot execution of REPIC with *𝒮*_init_ as REPIC(*𝒮*_init_),

#### Iterative mode

In the iterative mode, REPIC is used as described in Pseudocode 1. Here, *I* is the number of iterations chosen by the user. *M* is available cryo-EM micrographs split into cross-validation subsets (see Preprocessing below). *T*_*i*_ is the set of training labels (particle coordinates) used for *ab-initio* picker training in iteration *i*.

##### Pseudocode 1: REPIC — iterative mode

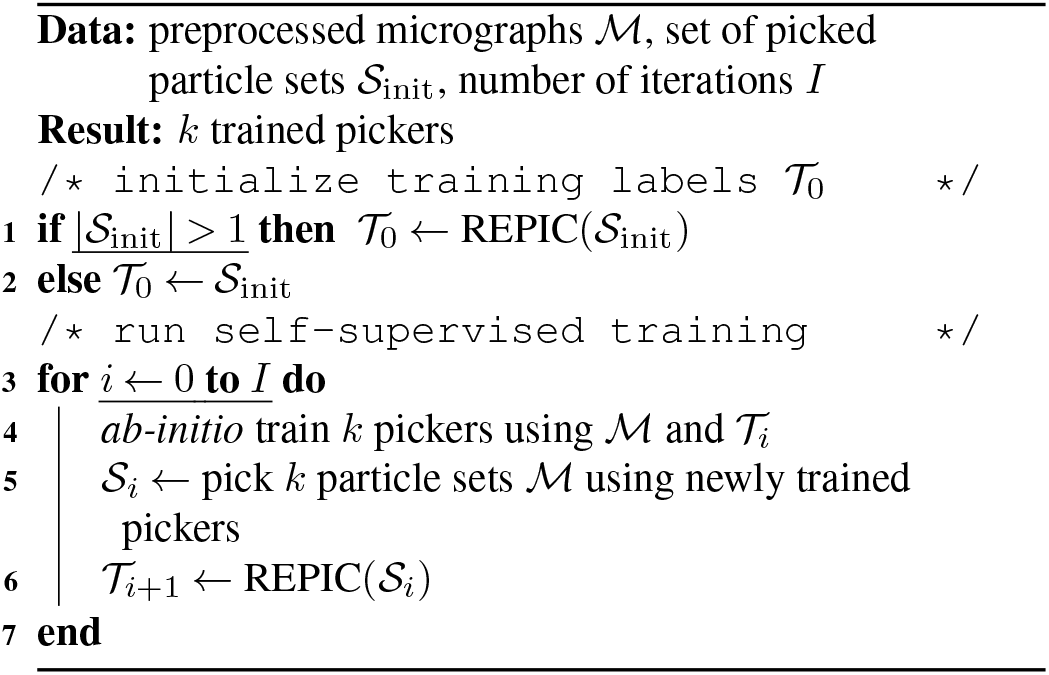

Cross-validation (training, validation, and testing) subsets *M* for Pseudocode 1 are created by sampling micrographs based on their mean defocus value. Mean defocus values are calculated from the output of a CTF estimation job in RELION v3.1.3 (45) using CTFFIND4 v4.1.14 (46). Specifically, ‘defocus 1’ and ‘defocus 2’ values are averaged per micrograph. Micrographs are then grouped into three bins (low, medium, and high) using their mean defocus values. Subsets are generated by randomly sampling (without replacement) three micrographs at a time, one from each bin. If defocus values are not available (e.g., EMPIAR-10057), all micrographs are randomly grouped into three equally sized bins. Training and validation sets are built first to ensure algorithms are exposed to the entire range of defocus values during picker training. For each dataset, validation subsets consist of six micrographs, and the remaining micrographs were initially split 20-80 between the training and testing subsets.

### Algorithm evaluation

For all datasets, picked and consensus particle sets are evaluated using published particle sets found on the EMPIAR resource. These published sets are used in place of a ground truth as the norm or expected picked particle set, which we refer to as the “normative”. Before evaluation, picker output was filtered for false positives using author-suggested thresholds (see Supplemental Information).

Precision and recall were calculated using micrograph pixels *P*, where *p*_*i*_ = 0 and *p*_*i*_ = 1 are a pixel found in the background region of a micrograph or a particle bounding box, respectively. A true positive (TP) is *p*_*i*_ =1 for both the normative and compared particle set. A false positive (FP) is *p*_*i*_ =0 in the normative and *p*_*i*_ =1 in the compared set. A false negative (FN) is *p*_*i*_ =1 in the normative but *p*_*i*_ =0 in the compared set. TPs, FPs, and FNs are summed over *P* before calculating either evaluation. Reported precision and recall values are computed from all testing micrographs in a dataset.

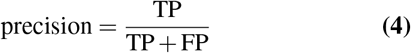

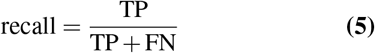

When available, the final particle set that produced a published density (e.g., EMPIAR-10454) is used as the normative particle set. If this final particle set is not available (e.g., EMPIAR-10057), the normative particle set consists of all particles found in the EMPIAR entry.

Initial analyses showed that the published final particle set of EMPIAR-10005 produced lower resolution reconstructions compared to the initial particle set. Missing summed frame micrographs in the EMPIAR entry reduced the final particle set from 35, 645 to 32, 387 particles. Therefore, the EMPIAR-10005 normative particle set was taken to be the published initial particle set reduced by the number of available micrographs (80, 443 particles - see Supplemental Data File 2).

### 3D reconstruction procedure

3D reconstruction was performed in RELION v3.1.3 (45). Soft masks were generated from published maps (see Dataset description in Online Methods) using a RELION mask generation job. For each particle set, a RELION 3D auto-refinement job was provided the corresponding soft mask, published density (low-pass filtered to 64 Å), and extracted particle images to produce a reconstruction. No particle filtering in RELION (either by 2D or 3D methodology) was performed on any particle set analyzed in this study. CTF correction was not performed for EMPIAR-10057 because it is an in-focus, phase-plate dataset. Final, unmasked reconstructions were generated using a RELION post-processing job. Unmasked normative reconstructions were then used to generate soft masks that were applied to their corresponding normative, consensus, and picker reconstructions (i.e., all reconstructions in the same row of a figure). RELION 3D auto-refinement jobs that had significantly longer runtimes (>24 hours) than the runtime for the normative particle set were aborted (e.g., DeepPicker EMPIAR-10005 reconstruction displayed in Figure 2 — see Supplemental Data File 3). For these particle sets (indicated by ‡ in figures), a single half map from the last-completed iteration of the RELION 3D auto-refinement job was used. Default RELION mask generation, 3D auto-refinement, and post-processing job parameters were used unless otherwise specified in Supplemental Data File 2.

### 3D reconstruction analysis

Masked reconstructions were registered to their corresponding normative reconstruction using UCSF Chimera (50) (https://www.cgl.ucsf.edu/chimera/ — ‘Fit in Map’ tool and ‘vop resample’ command) before Fourier shell correlation (FSC) calculation. Reconstructions resulting from either picker or consensus particle sets were compared to their corresponding normative map by calculating an FSC between both the masked and registered maps (a “cross-FSC”). A cross-FSC threshold of FSC= 0.5 was used as particle sets contributing to reconstructions may share particles and are not guaranteed to be independent. The reported cross-FSC resolution is the resolution where a map’s similarity to the normative map decreases below 50%. Half map FSCs are included as reference and their reported resolutions use the gold-standard FSC threshold (FSC= 0.143).

UCSF Chimera was used to visualize all maps. Normative maps were used to set the density threshold for maps resulting from either consensus or picker picked particle sets. The density threshold for half maps from aborted RELION 3D auto-refinement jobs (indicated by ‡ in figures) was set in an ad hoc manner to better visualize the obtained map.

## Data availability

Cryo-EM datasets used in this study are publicly available at the EMPIAR resource (https://www.ebi.ac.uk/empiar/): https://www.ebi.ac.uk/empiar/EMPIAR-10005/, https://www.ebi.ac.uk/empiar/EMPIAR-10017/, https://www.ebi.ac.uk/empiar/EMPIAR-10057/, and https://www.ebi.ac.uk/empiar/EMPIAR-10454/.

## Code availability

The source code for REPIC is available on GitHub at https://github.com/ccameron/REPIC. REPIC is licensed under the BSD-3-Clause.

A copy of the REPIC GitHub repository is available in the Gerstein lab’s GitHub account for posterity: https://github.com/gersteinlab

## ACKNOWLEDGEMENTS

Authors acknowledge financial support from the National Institute of Neurological Disorders and Stroke and National Institute of General Medical Sciences of the National Institutes of Health under award numbers R01NS021501 (F.J.S.) and R01GM125769 (H.D.T.), and the Albert L. Williams Professorship funds (M.B.G.). We would also like to thank Pengxin Chai, Swapnil Devarkar, Mihir Gowda, Jonathan Koss, Susanna Liu, Ran Meng, Daniela Nicastro, David Peng, Gryte Satas, and members of the Gerstein, Sigworth, and Tagare laboratories for their useful discussions during the development of this project and manuscript.

## AUTHOR CONTRIBUTIONS

C.J.F.C. conceptualized the study. C.J.F.C and H.D.T. were responsible for the methodology. C.J.F.C performed investigations. C.J.F.C and S.J.H.S. curated the data. C.J.F.C. performed visualizations. F.J.S., H.D.T., and M.B.G were responsible for funding acquisition. C.J.F.C. wrote the original manuscript draft. All authors reviewed and edited the manuscript.

## Notes

### Competing Interest Statement

The authors have declared no competing interest.

