## Supplemental Information for "REPIC — an ensemble learning methodology for cryo-EM particle picking"

**Supplemental Data File 1**
Randomly selected micrographs from the fatty acid synthase (FAS — EMPIAR-10454) dataset used in this study: https://github.com/ccameron/REPIC/blob/main/supp_data_files/supplemental_data_file_1.txt

**Supplemental Data File 2**
List of picker and RELION job parameters used in this study: https://github.com/ccameron/REPIC/blob/main/supp_data_files/supplemental_data_file_2.ods

**Supplemental Data File 3**
Tables describing the number of picked particles provided to RELION 3D auto-refinement jobs, and RELION 3D auto-refinement job runtimes and number of iterations: https://github.com/ccameron/REPIC/blob/main/supp_data_files/supplemental_data_file_3.ods

**Supplemental Methods**

**SPHIRE-crYOLO.** The SPHIRE-crYOLO particle picking software (29) was installed using Conda following author instructions: https://cryolo.readthedocs.io/en/stable/installation.html. A pre-trained SPHIRE-crYOLO picker (for low-pass filtered micrographs) was downloaded from: https://owncloud.gwdg.de/index.php/s/AdVdYdcCg4XaNRw. During iterative-mode REPIC, SPHIRE-crYOLO's low-pass micrograph filtering and pre-trained picker was used. To improve stability across iterations, the random number generator seed of SPHIRE-crYOLO was set to 1. Before evaluating SPHIRE-crYOLO picked particle sets, a particle score $s$ threshold of $s>0.3$ was used to filter false positives (author instructions).

**DeepPicker.** The DeepPicker particle picking software (19) was downloaded from the following GitHub repository: https://github.com/nejyeah/DeepPicker-python. A pre-trained DeepPicker picker is included in this GitHub repository. To improve DeepPicker particle picking efficiency, image preprocessing (specifically image resizing) was re-implemented in PyTorch (51). The resulting implementation is 10 times faster with a particle picking runtime of 3–5 seconds per micrograph. Minor additional software changes were made to ensure compatibility with REPIC's iterative mode (command line specification of validation micrographs, particle coordinates file paths, etc.). Instructions for installing and updating DeepPicker are available in the REPIC GitHub repository: https://github.com/ccameron/REPIC/blob/main/docs/deeppicker.md. Before evaluating DeepPicker picked particle sets, a threshold of $s>0.5$ was used to filter false positives (author instructions).

**Topaz.** The Topaz particle picking software (26) was installed using Conda following author instructions: https://github.com/tbepler/topaz. A pre-trained picker is included with the Conda installation. During micrograph preprocessing, all pixels were sampled to estimate the Gaussian mixture model's parameters. Topaz authors do note that the default sampling parameter (10% of pixels) introduces sampling error. In our experience, sampling all pixels did not significantly increase runtime but increased picker stability and performance. To obtain scores $s$, Topaz log-likelihood ratios $r$ of picked particles were transformed to probabilities before thresholding:

$$s = \frac{1}{1+\mathrm{e}^{-r}} \quad \textbf{(1)}$$

When training a Topaz picker during REPIC's iterative mode, the fraction of positive examples in a mini-batch (i.e., 'minibatch-balance') is set to the expected fraction of positive pixels. This expectation is the number of pixels found in particle detection boxes over the total number of pixels. During iterative-mode REPIC, consensus particles from the training set are used to calculate this fraction. For the initial round of picking, the default minibatch-balance value for Topaz is used. Before evaluating Topaz picked particle sets, a threshold of $s>0.5$ was used to filter false positives (author instructions).

2**D class average generation.** RELION v3.1.3 (45) 2D classification jobs were used to generate class averages for all particle sets. Default job parameters were used except for the number of classes (set to 32), the mask diameter (dataset specific — see Supplemental Data File 2), and limiting resolution of the expectation step (10 Å). Contrast transfer functions were ignored until the first peak. Note - 2D classes were not used to filter any particle sets used in this study

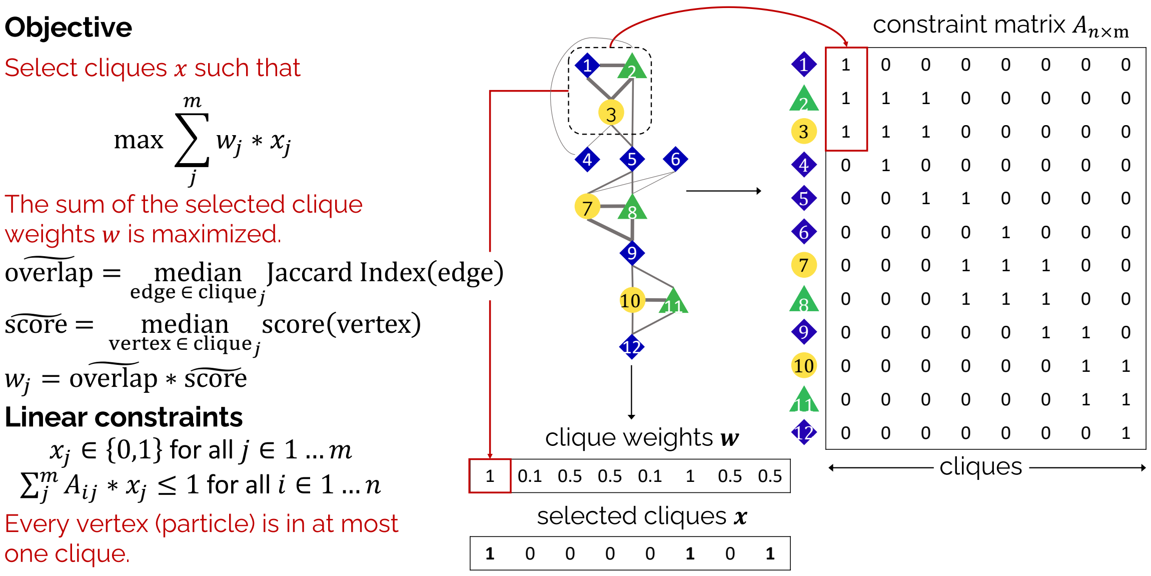


**Supplemental Figure S1 Integer linear programming framework**. Schematic of the integer linear programming (ILP) framework, where an optimal subset of cliques $x$ are selected (from the total number of cliques $m$) such that clique weights $w$ are maximized. Each clique weight $w_j$ is the product of the median overlap (measured by the Jaccard index) and median picker score of clique members. The linear constraints and constraint matrix $A$ ensure that cliques are binarily chosen and each particle can only be assigned to a single selected clique.

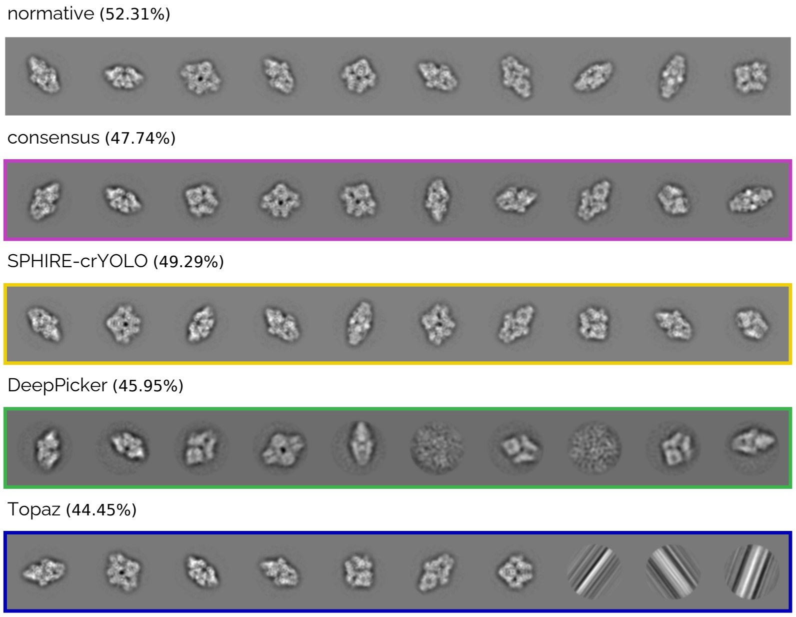


**Supplemental Figure S2 $\beta$-gal 2D classes resulting from out-of-the-box pickers**. Top ten of 32 RELION 2D classes of $\beta$-gal (EMPIAR-10017) resulting from normative (grey — *top*), consensus (purple), SPHIRE-crYOLO (yellow), DeepPicker (green), and Topaz (blue) picked particle sets. The total class distribution for the top-ten classes is listed in parentheses. The quality of RELION 2D classes (resolution and relevant views) for each particle set follows that of the downstream RELION 3D reconstructions (similar observations for other datasets). Note - 2D classes were not used to filter any particle sets used in this study

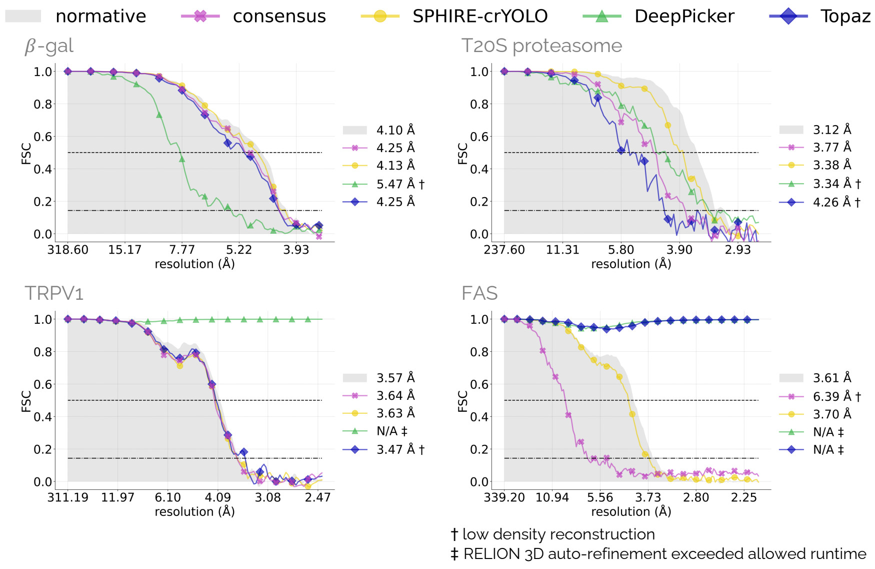


**Supplemental Figure S3 Half map FSCs for one-shot REPIC using out-of-the-box pickers**. Normative, out-of-the-box picker, and one-shot REPIC half map Fourier Shell Correlation (FSC) curves of RELION 3D auto-refinement half maps generated from $\beta$-gal (EMPIAR-10017); in-focus, phase-plate T20S proteasome (10057); TRPV1 (10005); and FAS (10454) datasets. Reported half map FSC resolutions are at FSC$= 0.143$.

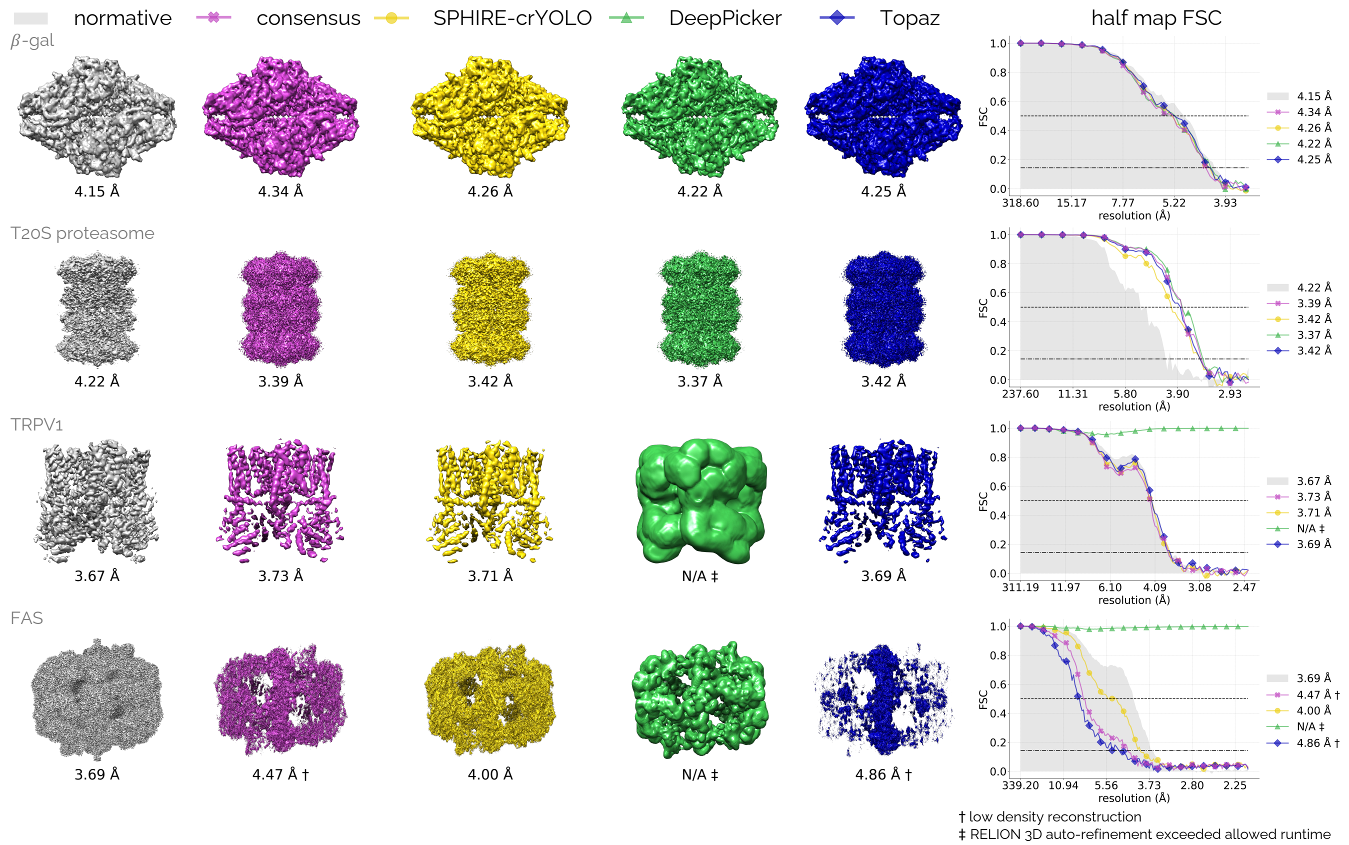


**Supplemental Figure S4 Final densities obtained from REPIC's iterative mode initialized using out-of-the-box pickers**. Normative (grey — *left*), consensus (purple), SPHIRE-crYOLO (yellow), DeepPicker (green), and Topaz (blue — *right*) reconstructions obtained from the final iteration. $\beta$-gal (EMPIAR-10017), T20S proteasome (10057), TRPV1 (10005), and FAS (10454) datasets were allowed 16, 24, 16, and 16 iterations of iterative-mode REPIC, respectively. Reported resolutions are calculated using an FSC$= 0.143$ and the half maps from the final iteration of a RELION 3D auto-refinement job. Reconstructions shown above are used to generate the cross-FSC curves in Figure 3C (*right* column).

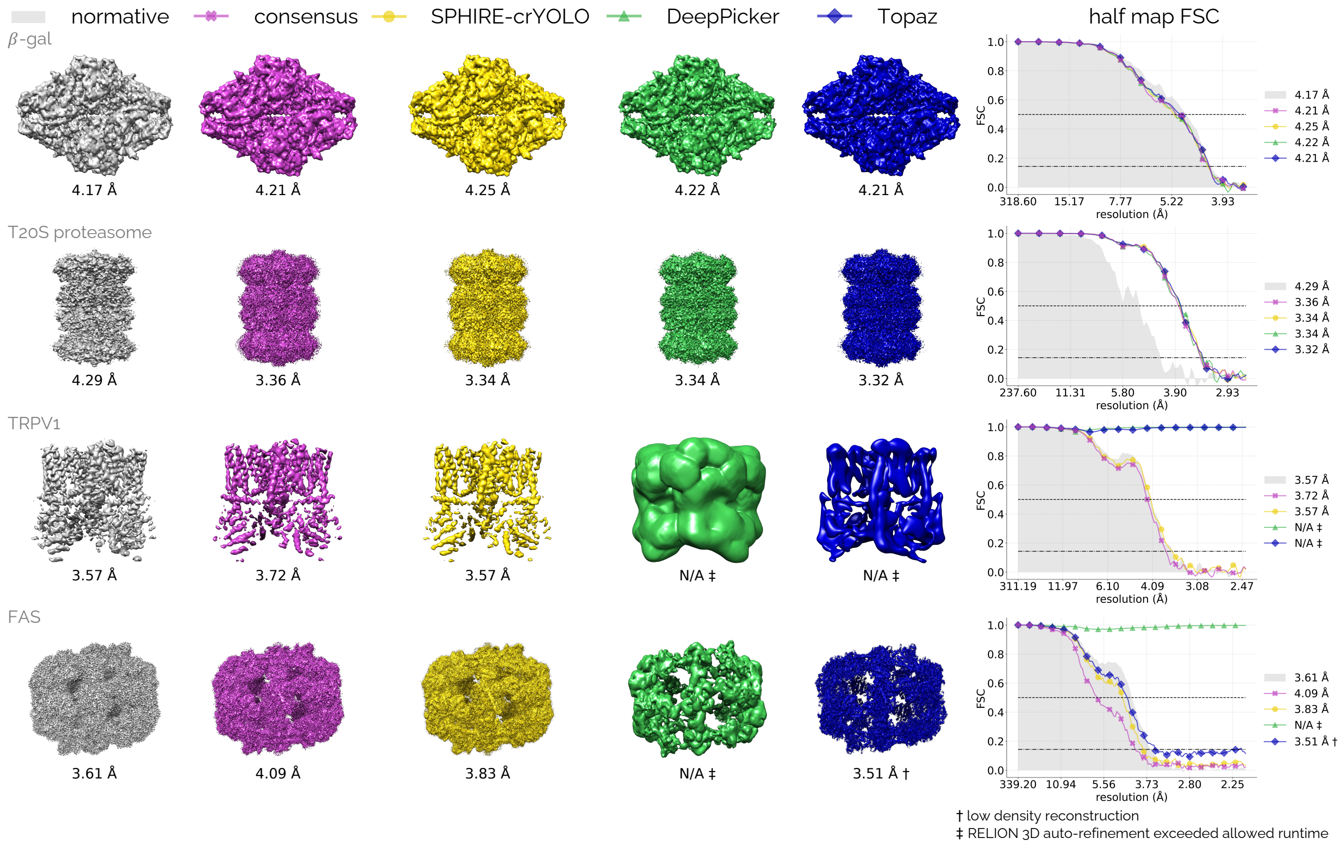


**Supplemental Figure S5  Final densities obtained from REPIC's iterative mode initialized with minimal manual picking**. Normative (grey — *left*), consensus (purple), SPHIRE-crYOLO (yellow), DeepPicker (green), and Topaz (blue — *right*) densities obtained from the final iteration. $\beta$-gal (EMPIAR-$10017$), T20S proteasome ($10057$), TRPV1 ($10005$), and FAS ($10454$) were allowed $16$ iterations of iterative-mode REPIC. Reported resolutions are calculated using an FSC$=0.143$ and the half maps from the final iteration of a RELION $3$D auto-refinement job. Reconstructions shown above are used to generate the cross-FSC curves in Figure 4C (*right* column).

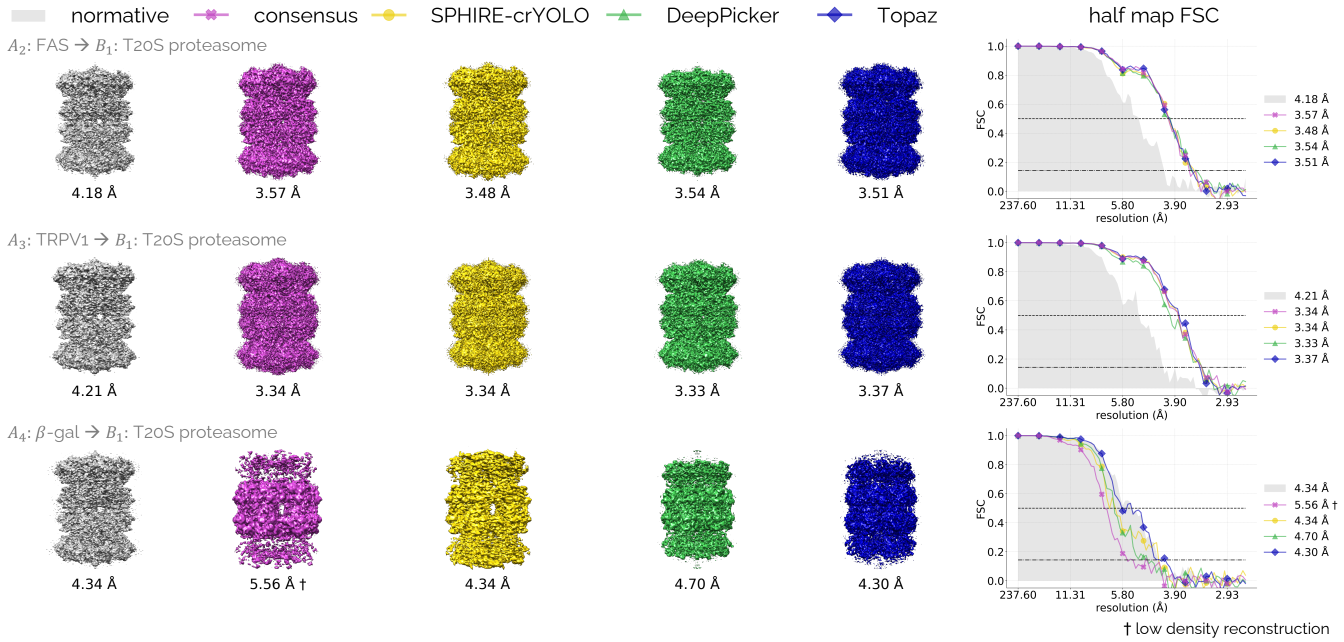


**Supplemental Figure S6  Final T20S proteasome densities obtained from *ab-initio* transfer learning using iterative-mode REPIC**. Normative (grey — *left*), consensus (purple), SPHIRE-crYOLO (yellow), DeepPicker (green), and Topaz (blue) T20S proteasome (EMPIAR-$10057$) reconstructions resulting from the final iterations of *ab-initio* transfer learning using pickers pre-trained on either FAS ($10454$ — *top*), TRPV1 ($10005$ — *middle*), and $\beta$-gal ($10017$ — *bottom*) datasets. Reported resolutions are calculated using an FSC$=0.143$ and the half maps from the final iteration of a RELION $3$D auto-refinement job. All instances of iterative-mode REPIC were allowed eight iterations to converge. FAS and TRPV1 pre-trained pickers lead to high-resolution reconstructions of T20S proteasome.

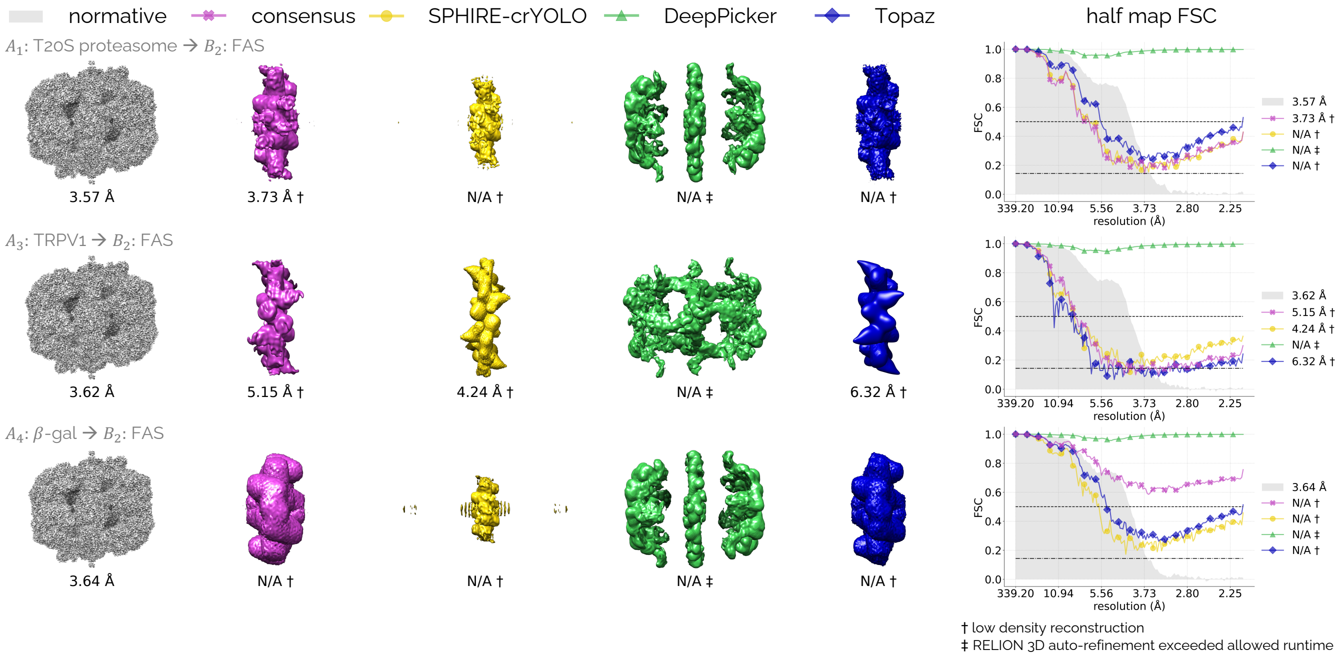


**Supplemental Figure S7 Final FAS densities obtained from *ab-initio* transfer learning using iterative-mode REPIC**. Normative (grey — *left*), consensus (purple), SPHIRE-crYOLO (yellow), DeepPicker (green), and Topaz (blue) FAS (EMPIAR-$10454$) reconstructions resulting from the final iteration of *ab-initio* transfer learning using pickers pre-trained on either T$20$S proteasome ($10057$ — *top*), TRPV$1$ ($10005$ — *middle*), and $\beta$-gal ($10017$ — *bottom*). Reported resolutions are calculated using an FSC$= 0.143$ and the half maps from the final iteration of a RELION $3$D auto-refinement job. All instances of iterative-mode REPIC were allowed eight iterations and failed to converge on a particle set that produced a high-resolution reconstruction.

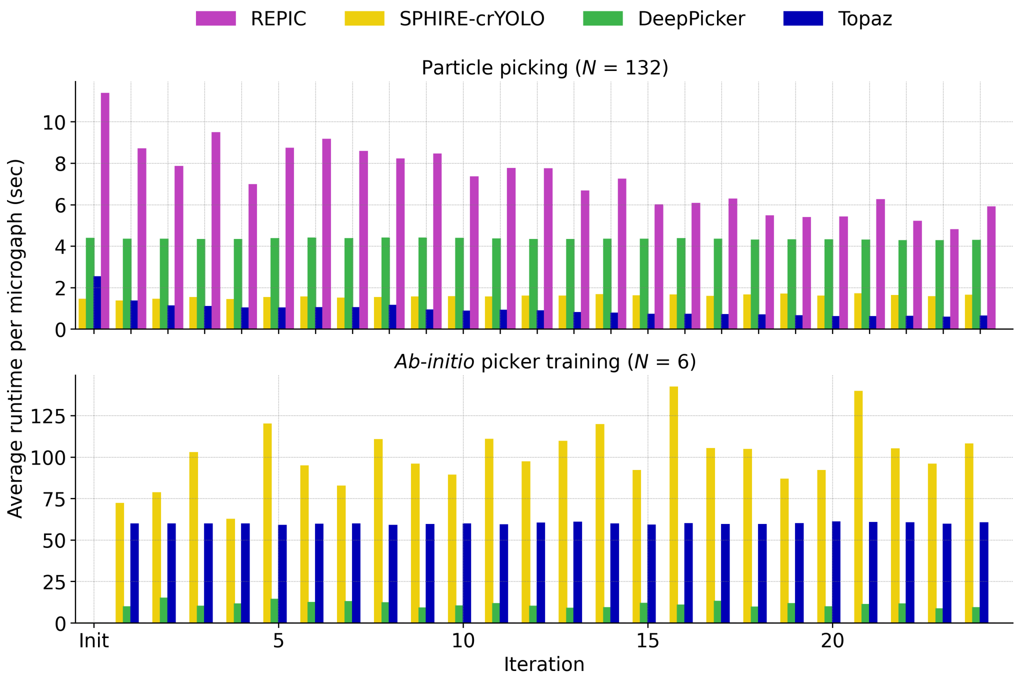


**Supplemental Figure S8 Algorithm runtimes**. Individual algorithm runtimes (in seconds per micrograph) for REPIC (purple), SPHIRE-crYOLO (yellow), DeepPicker (green), and Topaz (blue) for REPIC's iterative mode applied to the T$20$S proteasome dataset (EMPIAR-$10057$ — Figure 3B). $N$ is the number of micrographs used in either particle picking (*top*) or picker training (*bottom*).
